# Studies on Forecasting of Incidence of Leaf Mold in Tomato and Fungicide-spray Scheduling

**DOI:** 10.1101/2020.09.16.299503

**Authors:** Mun Haeng Lee

## Abstract

Tomato leaves were inoculated with 1×10^4^conidia·mL^−1^ and placed in a dew chamber for 0 to 18hrs at 10 to 25°C. Eighteen days after inoculation, leaf mold incidence appeared in 9hr treatment of leaf wet duration and the proper temperature was 15 to 20°C. In 10 and 25°C treatments, the incidence rate was 0% and 4.2%, respectively. The most important factors regard to leaf mold incidence were leaf wet duration and temperature. Considering the proper tomato growth temperature is 15-25°C, control by leaf wet duration is easier than control by temperature to prevent leaf mold incidence. After leaf mold inoculation, incidence appeared at 20 days and the latency period was estimated as 14-15 days. The leaf mold incidence rate was the highest at 15°C and 20°C. When ‘trihumin’ (triflumizole 30%) was applied at 12 hr of leaf wet duration, the control effect was the highest at 90% up to 168 hr but after 240 hr, it dropped to 60%. When ‘Demani B’ (polyoxin B 50%) was applied at 12 hour of leaf wet duration, the control effect was the highest at 91% up to 144hr but after 240hr, it dropped to 58%, similar to trihumin application treatment. When ‘belqute’ (Iminoctadine tris 40%) was applied at 12 hours of leaf wet duration, the control effect was the highest at 93% up to 144 hr but after 240 hr, it dropped to 65%.

## Introduction

With increasing public interest in health, demand for fresh vegetables is on the rise (USDA, 1992), and, to meet this increasing demand, the use of polyethylene mulching, chemical fertilizers, and synthetic pesticides is also increasing accordingly (NRCBA, 1989). In the Mid-Atlantic region of the US, a pest management model is used for tomato production, which aims to prevent diseases occurring in the aboveground parts of tomato (Sikora et al., 1994), and such models are gaining ground as a typical model applicable to the southeast region (Bauske et al., 1998). As guided by most pest control methods based on management models, an efficient approach is to spray pesticide from the date of planting until the harvest time at an interval of seven to ten days, but this method may lead to the waste of pesticide because pesticide spraying is often conducted regardless of weather conditions or whether the target disease concerned has occurred. Notably, in greenhouse cultivation environments, as typically seen in Korea, where cultivation is less affected by external environments, this approach may be considered inefficient.

Another approach is to use a forecasting model to predict the incidence of disease. FAST was developed to prevent early blight caused by Alternaria solani, and TOMCAST is a prediction model, derived from FAST, which is effective in preventing early blight (by Alternaria solani), septoria leaf spot (by Septoria lycopersici), and anthracnose fruit rot (by Colletotrichum coccodes) (Pitblado, 1988; 1992). These prediction models were found to reduce the number of pesticide spraying rounds compared to when existing pest management methods were applied (Gleason et al., 1995; MacNab et al., 1993; Sikora et al., 1994).

In Korea, research on plant disease prediction models started in 1947 with an investigation conducted by the Basic Research Department of the Central Agricultural Institute of Technology with regard to the scattering status of conidia of hyphomycetes and cochliobolus miyabeanus (Anonymous, 1947).

Fungal diseases that are known to occur in the aboveground parts of tomato include leaf mold (by Fulvia fulva), grey mold (by Botrytis cinerea), powdery mold, black leaf mold (by Pseudocercospora fuligena), early blight, and late blight (by Phytophthora infestance) (Korean Society of Plant Pathology, 2009).

In Korea, a model to predict the occurrence of late blight was developed by upgrading the moving graph method currently applied to potato cultivation in the US so as to reflect monthly average temperatures, 7-day cumulative precipitation, relative humidity, and 7-day average minimum temperatures. Also, based on the finding that the occurrence of late blight is closely related to the duration of leaf wetness, a dew condensation model applicable to potato crops was developed using temperature, relative humidity, and wind speed data and further used to develop a plant disease prediction model (Hahm et al., 1978; Hwang et al., 1996). Meanwhile, Ahn et al. (1998a,b) devised a moving average model capable of predicting the date of disease onset based on daytime average temperatures and the number of days of high humidity, thereby verifying that the progressive model of late blight fit a logit model.

Unlike in the US, most tomatoes are grown under greenhouses in Korea, and thus the effects of precipitation or wind speed are negligible. Also, the temperature and humidity can be adjusted. The present study intends to develop a prediction model that will enable crop producers to prevent an environment where disease can occur in the first place, for example, by controlling heating devices or opening up roof windows so that the humidity can be properly adjusted in a timely manner.

When condensation occurs on leaves, the developed model will help establishing and providing an appropriate pesticide spraying schedule to promote tomato cultivation.

## Materials and Methods

### Study for Prediction Model of Leaf Mold in Tomato

#### Species and Seedlings

Summer King (Vegetable Research Center, cherry tomato) was sown in gardening bed soil (nursery soil) contained in a 40-hole connection port, and tomato seedlings with five to six foliage leaves were used for testing.

#### Strains and Inoculation Methods

Fulvia fulva was collected from tomatoes grown in tomato farms in Buyeo, Chungcheongnam-do, and monospore isolation was conducted on this strain in a PDA medium containing streptomycin. Subsequently, the isolated strain was inoculated into the V8 medium and subject to UV radiation for 30 minutes at an interval of two hours over the period of 20 days to promote spore production. The spore concentration was measured using a hemocytometer coupled with an optical microscope. Sterile water was added to dilute the solution, and, as a result, a spore suspension with a concentration of 1×10^4^ ml^−1^ was prepared. The tomato seedlings grown in the 40-hole connection port were inoculated with 40 ml of the spore suspension in spray form. Before the inoculation process, it was ensured that all their leaves remained sufficiently wet.

#### Condensation on Leaves and Treatment Temperatures

The tomato seedlings inoculated with the spore suspension were assumed to be in a condensation state while staying in a wet phase at 10, 15, 20, 25, and 30°C for the time period ranging from zero to 24 hours.

#### On-farm Investigation of Occurrence of Leaf Mold

The investigation was conducted on ten farms located in Buyeo and Nonsan, Chungcheongnam-do, and the temperature and condensation time were measured using Watchdog 1450 (Spectrum Tech., USA) and Leaf Wetness (Spectrum Tech., USA).

#### Investigation of Occurrence of Leaf Mold and Relevant Statistics

The investigation was conducted 20 days after the inoculation, and the incidence rate was estimated by comparison with the control group where the spore suspension was not inoculated. Sigmaplot was employed as statistics software.

### Study of Leaf Mold Management Modeling

#### Species and Seedlings

Summer King (Research Institute in Buyeo County, cherry tomato) was sown in gardening bed soil (nursery soil) contained in a 40-hole connection port, and tomato seedlings with five to six foliage leaves were used for testing.

#### Strains and Inoculation Methods

Fulvia fulva was collected from tomatoes grown in tomato farms in Buyeo, Chungcheongnam-do, and monospore isolation was conducted on this strain in a PDA medium containing streptomycin. Subsequently, the isolated strain was inoculated into the V8 medium and subject to UV radiation for 30 minutes at an interval of two hours over the period of 20 days to promote spore production. The spore concentration was measured using a hemocytometer coupled with an optical microscope. Sterile water was added to dilute the solution, and, as a result, a spore suspension with a concentration of 1×104 conidia · ml-1 was prepared. The tomato seedlings grown in the 40-hole connection port were inoculated with 40 ml of the spore suspension in spray form. Before the inoculation process, it was ensured that all their leaves remained sufficiently wet. The inoculated tomato seedlings were treated in a dew chamber at 20°C for 12 hours and moved to a 20°C incubator.

Chemical Agent Treatment. 30% of trimidazole, 50% of polyoxin B, and 40% of iminoctadine tris (Belkut) were diluted 1,000 times to achieve the target concentration and applied to the tomato seedlings within the time period ranging from zero to 240 hours after the inoculation of spores. Also, the control group, i.e., untreated seedlings, was prepared for comparison.

#### Investigation of Occurrence of Leaf Mold and Relevant Statistics

The investigation was conducted 20 days after the inoculation, and Sigmaplot was employed as statistics software.

## Results and discussion

### Development of Leaf Mold Prediction Model

The test tomato seedlings were inoculated with the spore suspension of Fulvia fulva with a concentration of 1×104 conidia · ml-1 and subsequently subjected to the condensation process in a wet phase at 10-30°C for the time period ranging from zero to 24 hours. When measured during the period from Day 5 to Day 20 after the inoculation, leaf mold started to occur when the condensation time exceeded nine hours, and the optimum temperature for the incidence was determined to be 15-20°C (Fig. 1). At 10°C, the disease did not occur, and, at 25°C, the incidence rate still remained low, i.e., 4% even when the condensation time exceeded 16 hours (Fig. 1). Given that the most critical factors that determine the occurrence of leaf mold are the leaf condensation time and temperature, and that the optimum temperature for tomato cultivation is 15-25°C, adjusting the leaf condensation time, rather than adjusting the temperature, is considered effective in preventing leaf mold. These observations are in line with the results obtained by Nita et al. (2003) and Park et al. (1992) that the incidence rate of zonate leaf spot on strawberries increased with increasing leaf condensation time, and that it was possible to estimate the occurrence time of anthracnose and the appropriate time to take pest management measures based on the timing of dew formation on grapes.

**Fig. 1.**
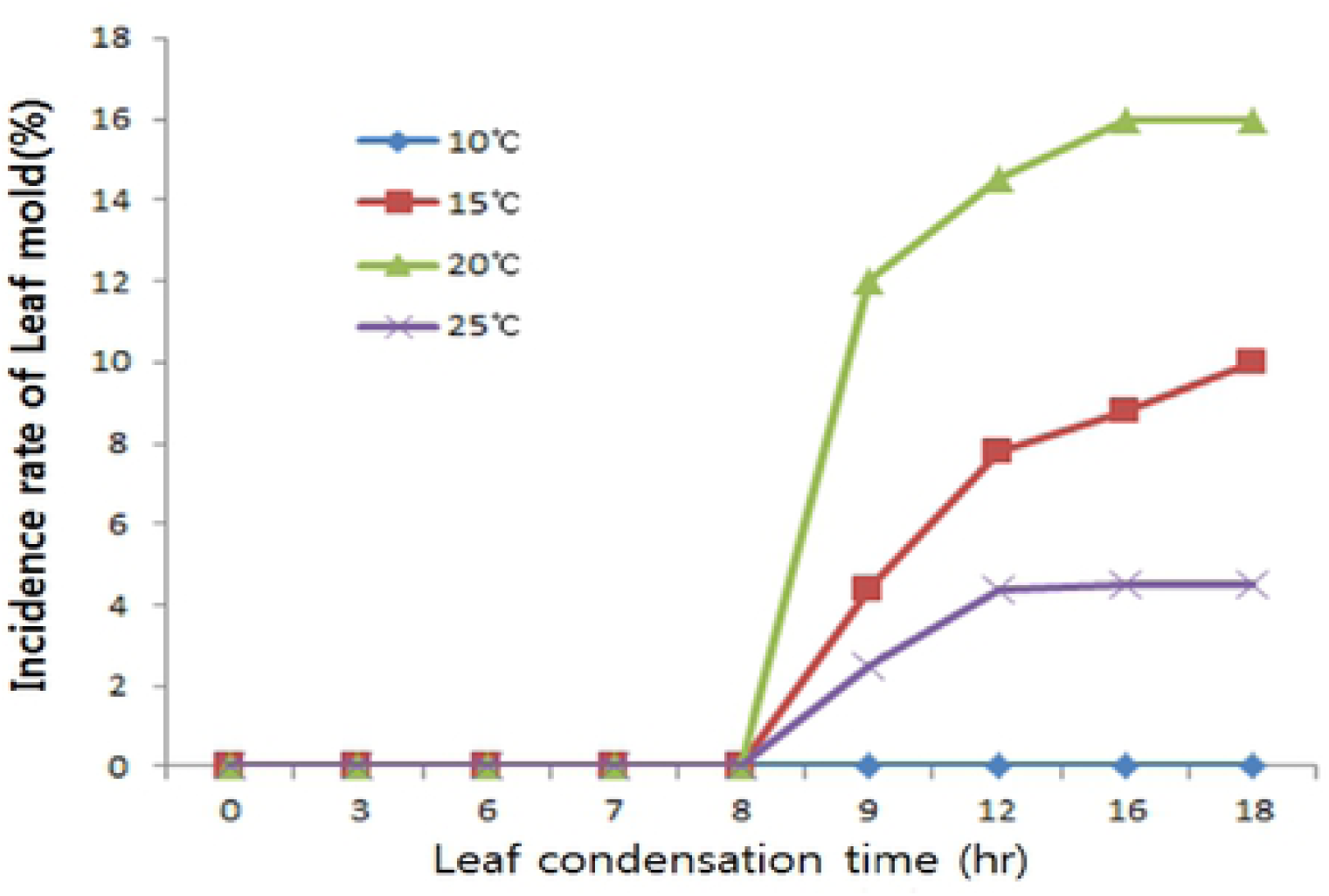
Effects of the condensation time of moisture and temperatures on the occurrence of leaf mold symptoms in the leaves of tomato plants

Given the findings above, when condensation on leaves occurs in a greenhouse, it is recommended to increase the temperature using a boiler and decrease the humidity by opening up side windows or roof windows. It is expected that these simple measures alone will sufficiently suppress the occurrence of leaf mold. Meanwhile, the tomato seedlings inoculated with leaf mold spores were monitored, and, as a result, the incidence rate of leaf mold was determined according to the temperature. The first incidence was observed 14 days after the inoculation, and thus the latency period is estimated to be 14 to 15 days (Table 1).

Fig. 1 Incidence rate of leaf mold, according to condensation time and temperature.

Table 1. Latency period of leaf mold according to temperature

### z Incidence rate of leaf mold, condensation time 12 hours

Ten farms in Buyeo and Nonsan, Chungcheongnam-do were investigated, and the results showed that the R-squared value between the leaf condensation time and the incidence rate of leaf mold was very high at 0.98, and the disease did not occur when the condensation time was less than seven hours (Fig. 2). These results are similar to those obtained by inducing the incidence of leaf mold using a wet phase (Fig. 1).

**Fig. 2.**
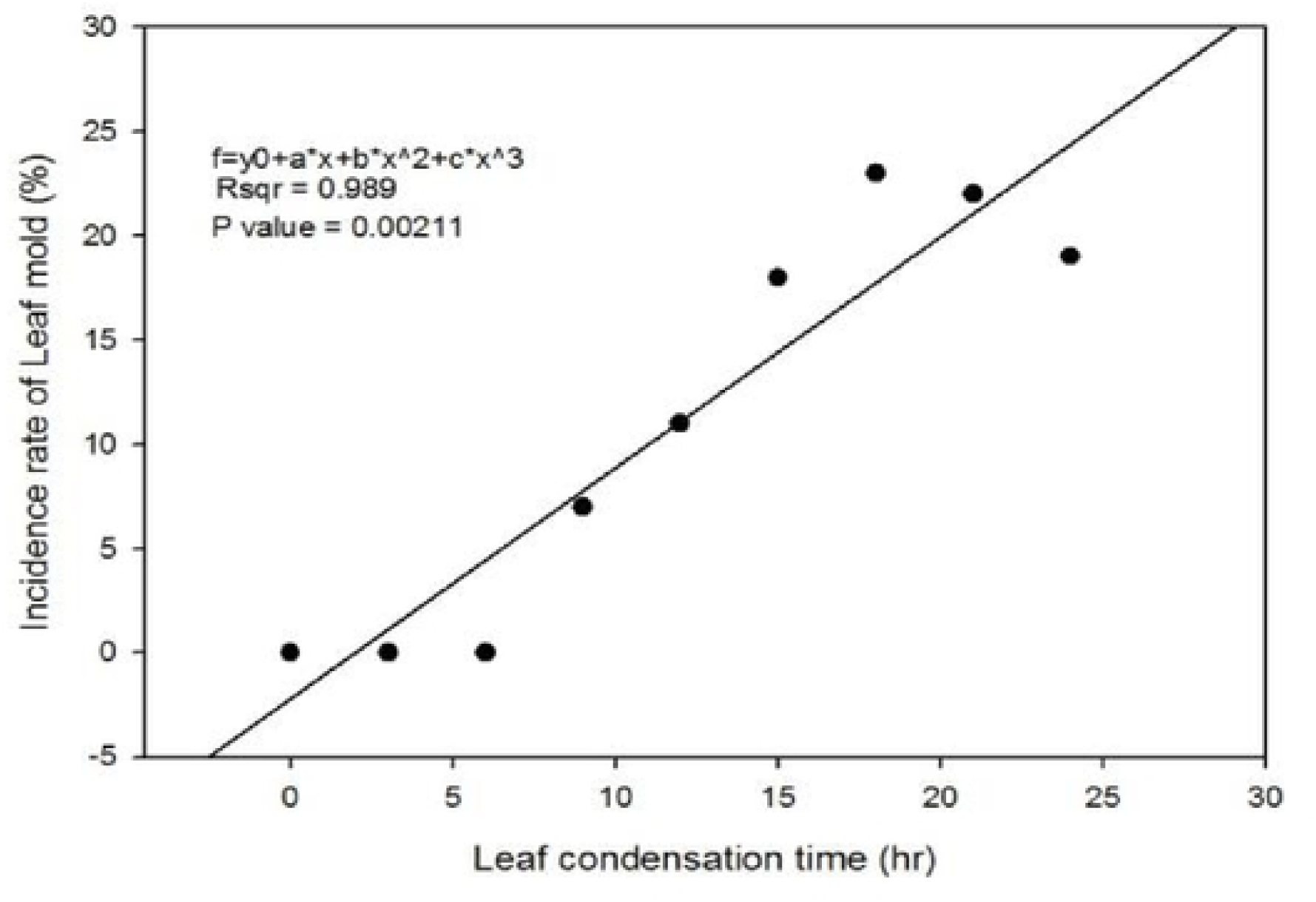
Comparison of the rate of leaf mold occurred according to the condensation time of moisture in the greenhouse of the farm

Fig. 2 Incidence rate of leaf mold, according to condensation time in farm field

### Study of Leaf Mold Management Modeling

Fig. 3 Control effect of leaf mold by chemical and treatment time after condensation time of 12 hour (A) triflumizole 30% × 2,985times; (B) polyoxin B 50% × 5,000times; (C) iminoctadine tris 40% × 2,000times.

**Fig. 3.**
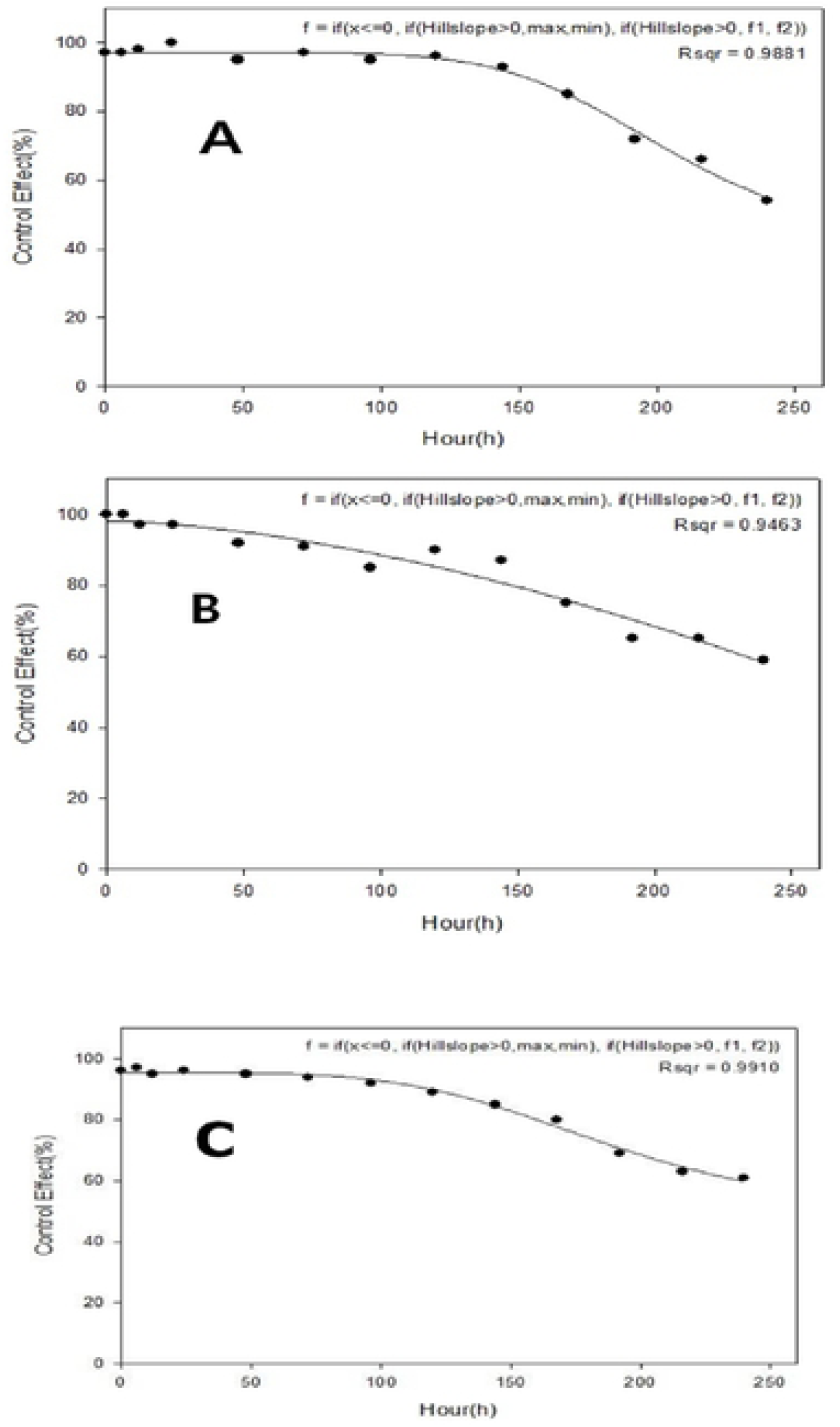
Control effect of leaf mold according to the treatment time of chemicals after 12 hrs of moisture condensation on the leaves (A) triflumizole 30%, 2,985times; (B) polyoxin B 50%, 5,000times; (C) iminoctadine tris 40%, 2,000times.

The tomato seedlings inoculated with Fulvia fulva spores were treated with chemical agents applied to leaf mold in tomato, i.e., trimidazole, polyoxin B, and iminoctadine tris (Belkut). After 12 hours of the condensation time, the incidence rate of leaf mold was investigated for 20 days (Fig. 3). When treated with trimidazole, polyoxin B, and iminoctadine tris 144 hours after the inoculation of spores, the control effects of these chemical agents were relatively high at 98, 88, and 86%, respectively, but when treated 240 hours after the inoculation, the figures decreased to 60, 60, and 65%, respectively. These observations show that the treatment time is more important than the type of chemical agent in preventing leaf mold. It should, however, be noted that the latency period of leaf mold is approximately 14 days (as shown in Table 1). Therefore, when it is found that leaf mold occurs on late leaves as a result of disease forecasting, it is necessary to remove those affected leaves and three earlier leaves suspected to be affected and immediately spray appropriate chemical agents so that the disease can be effectively contained. Park et al. (1992) reported that it was possible to reduce the amount of pesticide used to prevent grape anthracnose by determining the timing of pesticide spraying based on the incidence of condensation on grapes. Lee et al. (2017) reported that, similarly in addressing the incidence of tomato late blight, the earlier chemical agents are applied, the higher their control effects will be. The major findings of the present study indicate that it will be possible to reduce the total amount of pesticide used by taking pest management measures in a timely manner based on the leaf condensation time.

## Conclusions

Leaf mold did not occur at 10-30°C until the leaf condensation time reached eight hours. The first incidence was observed when the time reached nine hours. At an average temperature of 10°C, leaf mold did not occur, but the incidence rate increased to 4-10% at 15°C, 12-16% at 20°C, and 2.5-4.5% at 25°C, respectively. The latency period was determined to be 14 to 15 days at 20°C. To determine the control effects of chemical agents, tomato leaves were inoculated with a spore suspension with a concentration of 104/ml and subjected to the condensation process in a dew chamber at 20°C for 12 hours. The control effect of trimidazole (Trihumin) was 90% or higher when applied 168 hours after the spore inoculation and 60% when applied 240 hours after the inoculation. The control effect of polyoxin B (Demani B) was 90% or higher when applied 144 hours after the inoculation and 80% when applied 168 hours after the inoculation. The control effect of iminoctadine tris (Belkut) was 90% or higher when applied 144 hours after the inoculation but 80% and 65% when applied 168 hours and 240 hours after the inoculation, respectively. Based on these results, the optimum time to apply chemical agents was determined to be six to seven days after the infestation of pathogens. However, the latency period of leaf mold is approximately 15 days, and thus it is too late to effectively prevent the disease if chemical agents are applied after its incidence. Therefore, when it is found that leaf mold occurs on late leaves as a result of disease forecasting, it is necessary to remove not only those affected leaves but also two or three earlier leaves suspected to be affected so that the disease can be effectively contained. The forecasting model developed in the present study will help in determining under what conditions leaf mold occurs. Based on these predictions, crop producers can prevent such an environment where the disease can occur in the first place, for example, by running heating devices or opening roof windows and side windows. Also, if the incidence of the disease is expected, the timing of pesticide spraying can be properly adjusted and optimized according to the pest management schedule determined accordingly. All these measures together will sufficiently suppress the occurrence of leaf mold, thereby contributing to the stable cultivation of tomatoes.

## Acknowledgement

This study was carried out with the support of “Research Program for Agricultural Science &Technology Development(Project No. PJ0138912019) Administration” National Institute of Agricultural Science, Rural Development Administration, Republic of Korea.

## References

1. Ahn, J. H., Hahm, Y. I. and Park, E. W. 1998a. Development of ‘moving average method’ for prediction of initial appearance of potato late blight. Korean J. Plant Pathol. 14(1): 34–40.

2. Ahn, J. H., Hahm, Y. I. and Shin, K. Y. 1998b. Modeling for prediction of potato late blight(Phytophthora infestans) progress. Korean J. Plant Pathol. 14(4): 331–338.

3. Anonymous, 1947. Investigation if air born conidia dispersal of rice blast and Helminthosporium leaf spot pathogens in day and night time. Central Agricultural Technology Institute. An. Research Rept. 314

4. Bauske, E. M., Zehnder, G. M., Sikora, E. J., and Kemble, J. 1998. Southeastern tomato growers adopt intergrated pest management. HortTechnology. 8:50–44.

5. Bierhuizen, J. F. and Wagenvoort, W. A. 1974. Some aspects of seed germination in vegetables, 1. The determination and application of hit sums and minimum temperature for germination. Scientia Horticulture 2: 213–219.

6. Gleason, M. L., MacNab, A. A., Pitblado, R. E., Ricker, M. D., East, D. A., and Latin, R. X. 1995. Disease warning systems for processing tomatoes in eastern North America: Are we there yet? Plant Dis. 79:112–121.

7. Hahm Y. I., Hahn B. H. and J. D. Franckowiak. 1978. Forecasting Late Blight of Potatoes at the Alpine Area in Korea. Korean J. Plant. Prot. 17(2) : 81–87.

8. Hwang, B. S., Yun, J. I. and Lee, K. H. 1996. Using hourly weather data to determine dew periods of potato crops. Korean J. Plant. Prot. 12(4) : 445–452.

9. Korean society of plant pathology. 2009. List of plant diseases in Korea. 151–157.

10. Lee, M. H., Cho, S. K., Kim, Y. S., and Lee, D. J. 2017. Study of fordxasting and Scheduling gor fungicide sprays to control of Late Blight in Tomato. Kor. Soc. Int. Agric., 29(4) 454–458.

11. MacNab, A. A., and Gardner, R. G. 1993. Early blight defolination on tomatoes associated with cultivar and fungicide treatments, 1992. Biol. Cult. Tests 8:54.

12. National Research Council, Board on Agriculture(NRCBA). 1989. Alternative agriculture. National Academiy of Science, Washington, D. C. pp 195–244.

13. Nelson, S.C. 2008. Plant Disease-Late Blight of Tomato(Phytophthora infestance). Cooperative Extension Service. University of Hawai. www.ctahr.hawaii.edu/oc/freepubs/pdf/PD-45.pdf

14. Nita, M., Ellis, M. A. and L. V. Madden. 2003. Effects of Temperature, Wetness Duration, and Leaflet Age on Infection of Strawberry Foliage by Phomopsis obscurans. Plant Dis. 87:5 579–584

15. Park, E. W., Hur, J. S., and Yun, S. C. 1992. A Forecastig system for scheduling fungicide sprays to control grape ripe rot caused by Colletotrichum gloeosporioides. Korea J. Plant Path. 8:3 177–184.

16. Pitblado, R. E. 1988. Development of a weather-timed fungicide spray program for field tomatoes. Can J. Plant Pathol. 10:371.

17. Pitblado, R. E. 1992. The development and Implementation of TOM-CAST, a Weather Timed Fungicide Spray Program for Field Tomatoes. Ministry of Agriculture and Food, Ridgetown College of Agricultural Technology, Ridgetown Ontario, Canada.

18. Sikora, E. J., Bauske, E. M., Zehnder, G. W., and Hollingsworth, M. H. 1994. Evaluation of low-input fungicide spray programs for control of early blight on tomatoes. Highlights of Agricultural Research. Ala. Agric. Exp. Stn. 41:15.

19. United States Department of Agriculture(USDA), Economic Research Service. 1992. Americans are eating more fruits and vegetables. farmline. 13(7):8–13.

20. Yun, S. C., and E. W. Park. 1990. Effects of Temperature and Wetness Period on Infection of Grape by Colletrotrichum gloeosporioides. Korean J. Plant Pathol. 6(2): 219–228.

